# The primary structural photoresponse of phytochrome proteins captured by a femtosecond X-ray laser

**DOI:** 10.1101/789305

**Authors:** Elin Claesson, Weixiao Yuan Wahlgren, Heikki Takala, Suraj Pandey, Leticia Castillon, Valentyna Kuznetsova, Léocadie Henry, Melissa Carrillo, Matthijs Panman, Joachim Kübel, Rahul Nanekar, Linnéa Isaksson, Amke Nimmrich, Andrea Cellini, Dmitry Morozov, Michał Maj, Moona Kurttila, Robert Bosman, Eriko Nango, Rie Tanaka, Tomoyuki Tanaka, Luo Fangjia, So Iwata, Shigeki Owada, Keith Moffat, Gerrit Groenhof, Emina A. Stojković, Janne A. Ihalainen, Marius Schmidt, Sebastian Westenhoff

## Abstract

Phytochrome proteins control the growth, reproduction, and photosynthesis of plants, fungi, and bacteria. Light is detected by a bilin cofactor, but it remains elusive how this leads to activation of the protein through structural changes. We present serial femtosecond X-ray crystallographic data of the chromophore-binding domains of a bacterial phytochrome at delay times of 1 ps and 10 ps after photoexcitation. The structures reveal a twist of the D-ring, which lead to partial detachment of the chromophore from the protein. Unexpectedly, the conserved so-called pyrrole water is photodissociated from the chromophore, concomitant with movement of the A-ring and a key signalling aspartate. The changes are wired together by ultrafast backbone and water movements around the chromophore, channeling them into signal transduction towards the output domains. We suggest that the water dissociation is key to the phytochrome photoresponse, explaining the earliest steps of how plants, fungi and bacteria sense red light.

## Introduction

Discovered in 1959, phytochrome photosensor proteins are crucial for the optimal development of all vegetation on Earth (Butler et al. 1959; Gan et al. 2014; Quail et al. 1995). Prototypical phytochromes can exist in two photochemical states with differential cellular signalling activity, called red light-absorbing (Pr) and far-red light-absorbing (Pfr) state (Supplementary Fig. 1). As a result, phytochromes can distinguish two colours of light, providing plants, fungi, and bacteria with primitive two-colour vision. Light is detected by a bilin chromophore, which is covalently linked to the photosensory core of the protein (Wagner, Brunzelle, et al. 2005), comprising of PAS (Per/Arndt/Sim), GAF (cGMP phosphodiesterase/adenyl cyclase/FhlA) and PHY (phytochrome-specific) domains. Two propionate side chains additionally anchor it non-covalently to the protein (Fig. 1b). The signalling sites of the phytochrome are found in its C- and N-terminal output domains, which vary between species. Important for signalling is a stretch of amino acids in the PHY domain, called the PHY-tongue, which changes from a *β*-sheet in Pr into an *α*-helix in Pfr state (Essen et al. 2008; Takala et al. 2014; X. Yang et al. 2008). The chromophore connects to the PHY-tongue via a strictly conserved aspartatic acid, which is expected to play a crucial role in signal transduction.

**Figure 1:**
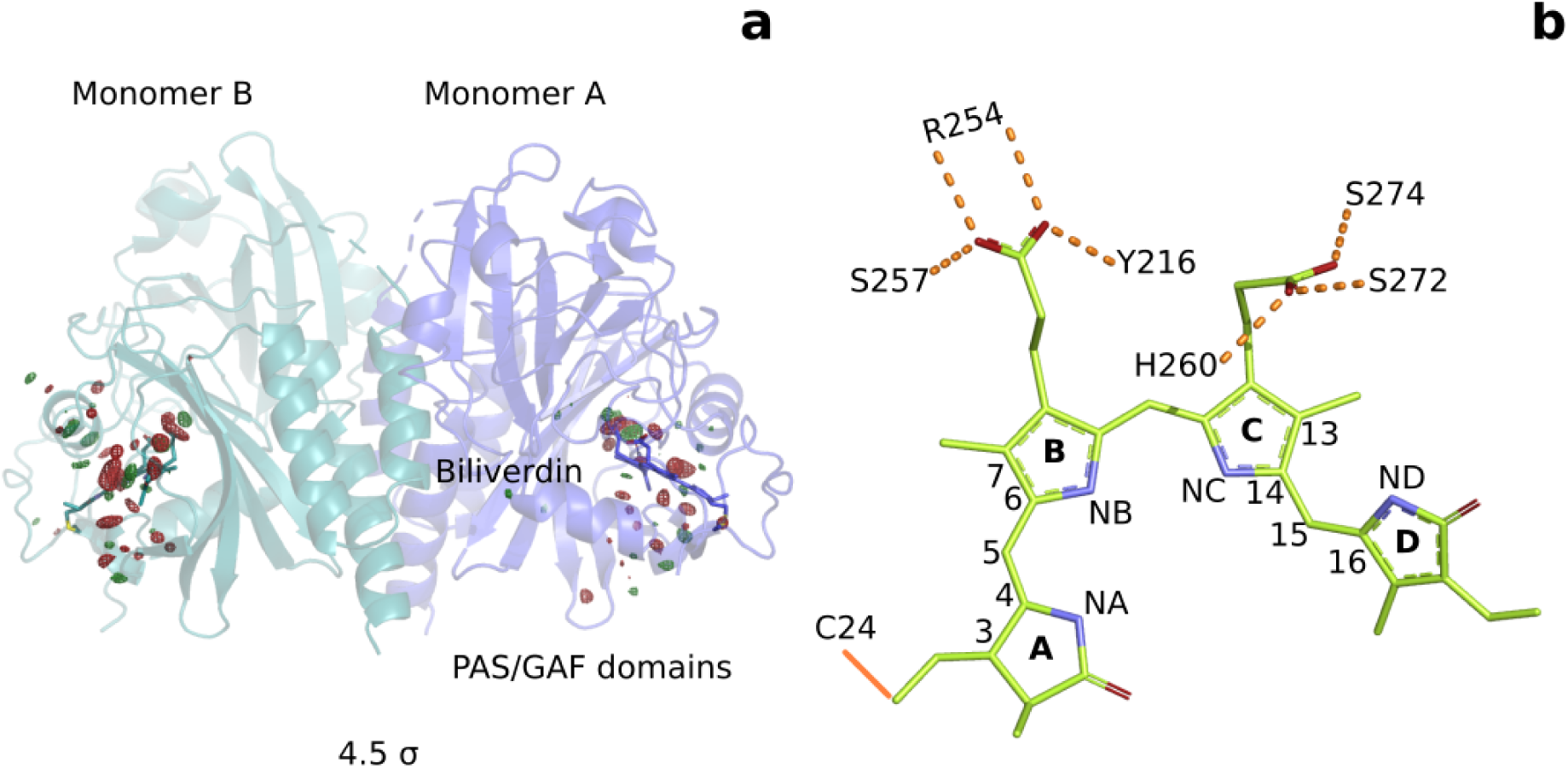
Photoinduced observed difference electron density features are focused on the chromophore binding pocket. (**a**). The observed difference electron density map at 1 ps is displayed together with the *Dr*BphP_*dark*_ structure. Red and green electron density peaks, contoured at 4.5 *σ*, denote negative and positive densities, respectively. Monomer A is coloured blue and monomer B is in aqua. (**b**). Schematic illustration of the biliverdin chromophore. The hydrogen-bonding networks between the propionate groups and the protein are marked with dashed lines. In *Dr*BphP, the chromophore is covalently linked to a cysteine residue in the PAS domain (solid line).

Key to phytochrome function is the primary photoresponse on picosecond time scales. Here, light signals are translated into conformational changes. The changes arise in the electronically excited bilin, but must then be transduced to the surrounding protein residues. This prepares the protein for formation of the first intermediate (Lumi-R for prototypical phytochromes), in which isomerisation of the D-ring has likely occurred (Dasgupta et al. 2009; Ihalainen et al. 2018; Rockwell et al. 2009; Y. Yang et al. 2012). The conformational changes are currently not well understood, because crystallographic observations of phytochromes directly after photoexcitation have not been available.

## Results

To address this gap of knowledge, we recorded time-resolved serial femtosecond X-ray crystallographic (SFX) data of the PAS-GAF domains of the phytochrome from *Deinococcus radiodurans* (*Dr*BphP) at 1 ps and 10 ps after femtosecond optical excitation. The experiments were performed in Japan, using the SPring-8 Angstrom Compact Free Electron Laser (SACLA) tuned to 7 KeV (Tono et al. 2015). For homogeneous excitation of the crystals, we photoexcited micrometer-sized crystals in a grease jet with a photon density of 1.9 mJ/mm^2^ (1*/e*^2^ measure) into the flank of the absorption peak at 640 nm (Supplementary Fig. 1 and 2). Our fluence dependent SFX data indicated that excitation density was in the one-photon regime, because lowering the excitation density resulted in a joint reduction of all difference signals (Supplementary Fig. 3). The refined structure in dark (*Dr*BphP_*dark*_, 2.07 Å resolution, (Supplementary Table. 1) was very similar to our previous dark structure solved by SFX (5K5B, RMSD 0.646 Å ^2^ and 0.610 Å ^2^ for monomers A and B) (Edlund et al. 2016), but the present crystals contained two monomers in the asymmetric unit.

From the time-resolved data, we calculated isomorphous difference electron density maps (*|F*_*o*_*|*^1*ps*^−*|F*_*o*_*|*^*dark*^), which report on the change of structure due to optical excitation. The map at 1 ps indicates many significant changes in difference electron density (Fig. 1a) above the background level of 3.0 standard deviations (*σ*) (Supplementary Fig. 4). The changes cluster around the chromophore, with the strongest negative densities for the pyrrole water (monomer A: −8.2*σ*, B: −9.4*σ*, Supplementary Table. 2). The map at 10 ps contains similar significant features, but at weaker overall intensity (pyrrole water A: −5.0*σ*, B:-6.5*σ*) (Supplementary Fig. 4). Monomer A has a lower signal strength than monomer B, but provided a clearer difference map around the chromophore. We focus our discussion on monomer A and the 1 ps time point, although all conclusions are supported by the features observed at 10 ps and monomer B (Supplementary Table. 2).

First, we inspect the D-ring region at 1 ps (Fig. 2). We observe strong negative difference density features on the atoms of the D-ring (marked **I, II, III**), correlating with density gains at both faces of the ring (**IV, V, VI**). These features strongly indicate that the D-ring twists. The positive feature **IV** homes the N-H and C=O groups, whereas **V** and **VI** indicate densities for the methyl and vinyl groups in the twisted ring (Fig. 2c). We refined a structural model (*Dr*BphP_1*ps*_) using extrapolated structure factors (Supplementary Fig. 5) (Pande et al. 2016). Excellent agreement was obtained between observed and calculated 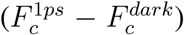 structurefactors, when the D-ring twists (C14-C15-C16-ND) from 25.1° and 20.2° (monomer A and B) in the dark to 58.2° and 89.4° at 1 ps (Fig. 2c and e).

**Figure 2:**
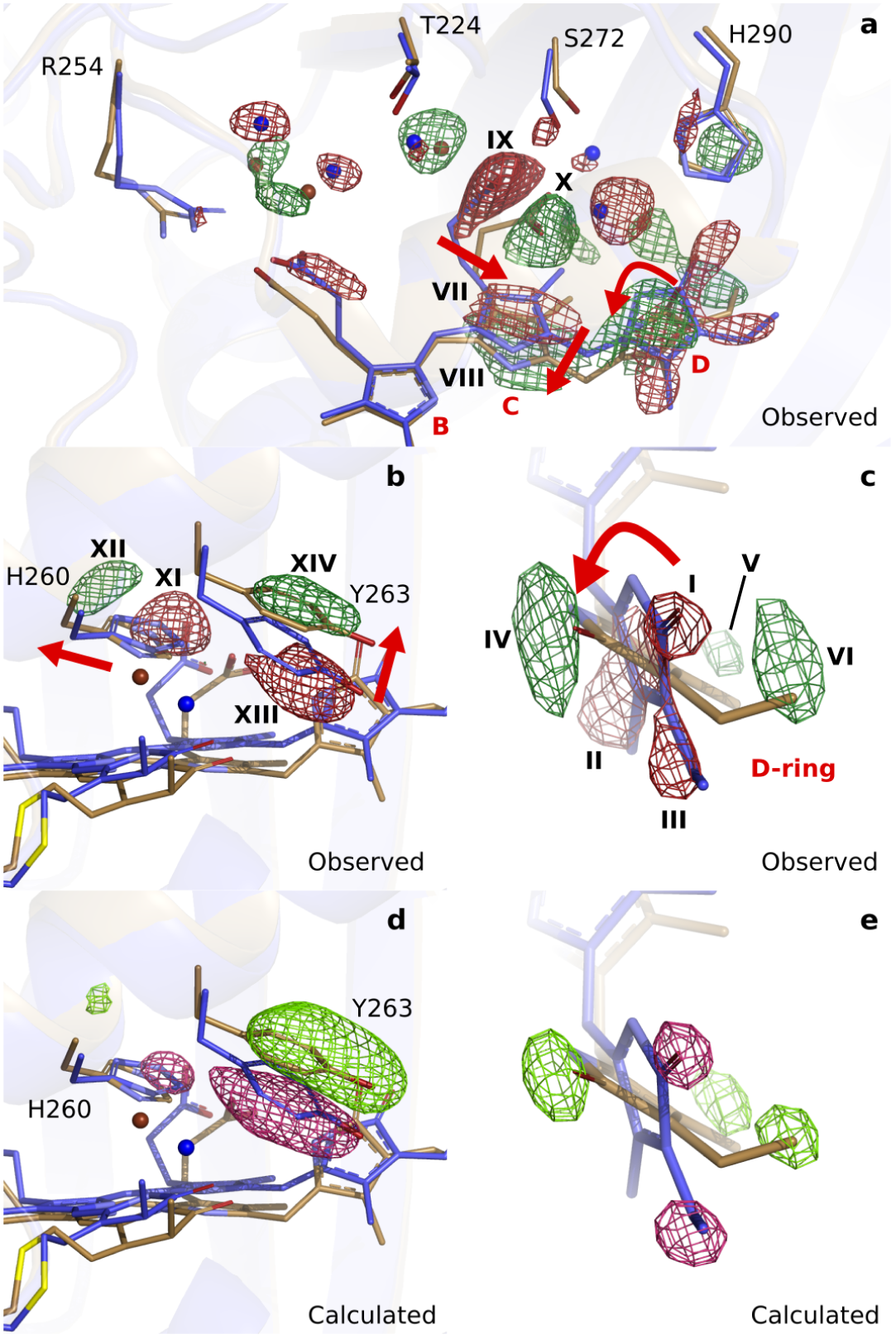
Observed and calculated difference electron densities reveal a twist of the D-ring and significant protein rearrangements. The observed difference electron density with the refined *Dr*BphP_*dark*_(blue) and *Dr*BphP_1*ps*_(beige) structures, shown for (**a**) the B-, C-, and D-ring surroundings, (**b**) the strictly conserved His260 and Tyr263, and (**c**) the D-ring. The calculated difference electron density shown for (**d**) His260 and Tyr263 and (**e**) the D-ring. The D-ring twists counter-clockwise when viewed along C15-C16 bond towards the C-ring. The observed difference electron density is contoured at 3.3 *σ*. And the calculated difference electron density is contoured at 3.5 and 5.0 *σ* for panel **d** and **e**, respectively. All the data correspond to monomer A.

Concomitant with the twist of the D-ring, the C-ring translates by approximately 0.69 Å as indicated by the correlated negative (**VII**) and positive (**VIII**) electron density (Fig. 2a). Furthermore, the C-ring propionate chain detaches from its conserved anchoring residues Ser272 and Ser274 (**IX** and **X**, Fig. 2a). The strictly conserved His260 retracts from its position (**XI** and **XII**) and Tyr263 moves upward at 1 ps (**XIII** and **XIV**, Fig. 2b). The water network connecting the C-ring propionate, the D-ring C=O, and His290 rearranges accordingly (Fig. 2a). The excellent agreement between calculated and observed difference maps confirms these observations (Fig. 2d and e, Supplementary Fig. 6). We conclude that the twist of the D-ring causes detachment of the C-ring propionate from the protein scaffold by dislocation of the C-ring, facilitated by the associated hydrogen bonding network.

Turning our attention to the B-ring, we find that the B-ring propionate breaks its salt bridge to Arg254 (Supplementary Fig. 7). However, this is not caused by movements of the chromophore backbone as we observe little change on the B-ring itself. Instead, we find that a water bridge between the B- and C-ring propionates is broken as indicated by negative difference electron densities on the waters (Supplementary Fig. 7). Additionally, the highly conserved helix from Ser257 to Val269, moves away from the chromophore by an average of 0.36 Å (monomer A) and 0.62 Å (monomer B) (distances relative to the pyrrole water, Supplementary Fig. 8). The changes of the D-ring are transduced to Ser257 via the side chains of His260 and Tyr263 and as a result the hydrogen bond of Ser257 to the B-ring propionate group breaks. The amino acids in the stretch are over 50% conserved (Supplementary Fig. 8), suggesting that it has evolved to transfer an ultrafast signal. We conclude that relaxation of the protein is necessary for the detachment of the B-ring propionate from the protein scaffold.

Based on spectroscopic data, it has been proposed that the D-ring of the bilin chromophore isomerizes around the C15-C16 bond from (Z) in Pr to (E) in the Lumi-R state (Dasgupta et al. 2009; Ihalainen et al. 2018; Rockwell et al. 2009; Song et al. 2014). Within the same model the D-ring is twisted in the excited state (Dasgupta et al. 2009). Our crystallographic data confirm the spectroscopic proposals and reveal that this leads to extensive and coordinated structural changes in the binding pocket, culminating in the detachment of the propionates from their protein anchors (Fig. 2). The result is a liberation of the chromophore from the binding pocket, which we propose to be necessary for large conformational rearrangements to occur in the downstream photoconversion to Pfr.

Next to the changes around the D-ring, the maps reveal strong difference electron density on the A-ring (**XVIII** and **XIX**), Asp207 (**XX** to **XXIII**) (Fig. 3a) and the pyrrole water (**XV**) (Fig. 3b). When interpreted and modelled as downward movement of the A-ring and Asp207 and photodissociation of the pyrrole water from the chromophore, excellent agreement between calculated and observed difference electron density is obtained (Supplementary Fig. 9). The A-ring is covalently attached to the protein backbone in phytochromes (Song et al. 2014), which renders complete isomerisation impossible, but is sufficiently flexible to accommodate the proposed changes. The pyrrole water may either move to feature (**XVI**), or occupy an anisotropic, worm-shaped feature which extends from the A-ring to the D-ring (**XVII**) (Fig. 3b).

**Figure 3:**
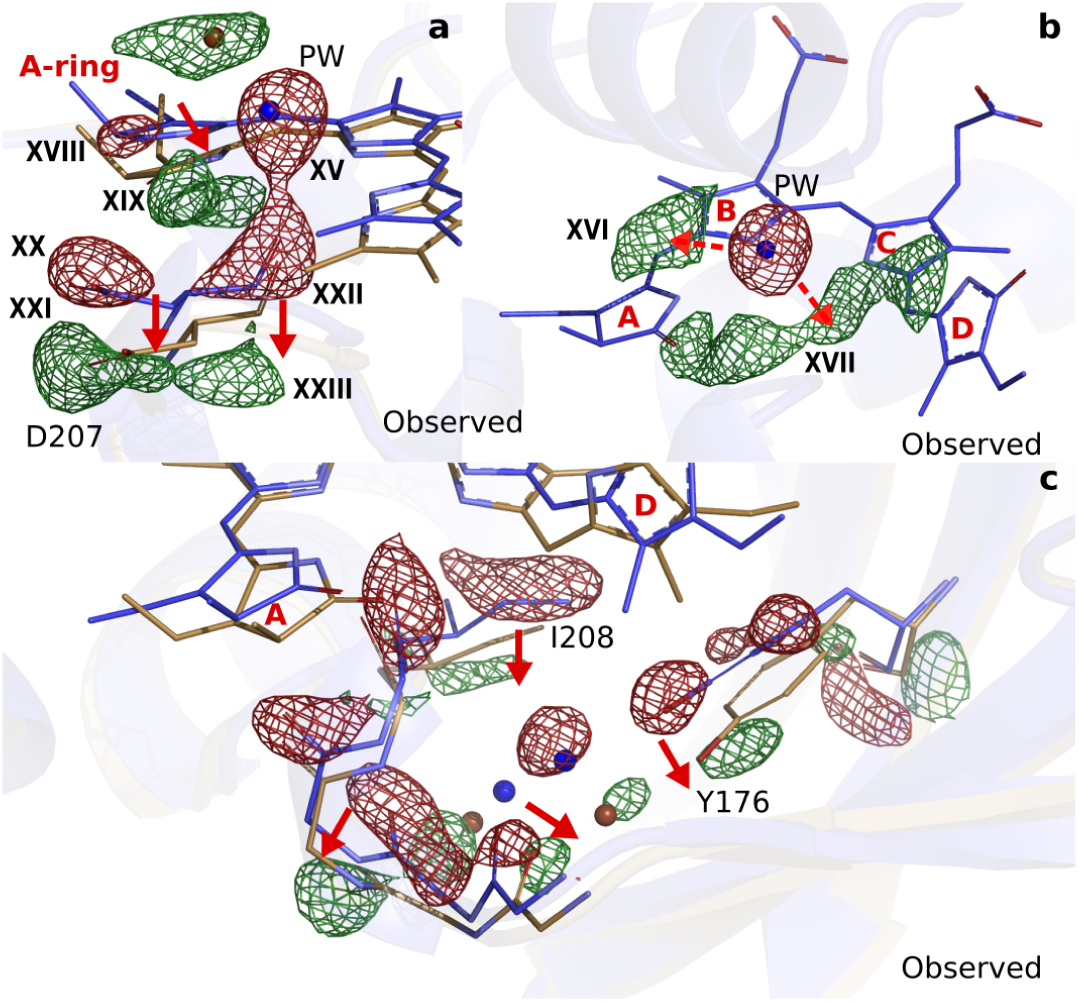
Photodissociation of the pyrrole water, displacement of the A-ring and its effect on the proteins scaffold. (**a**). The observed difference electron density displayed with the *Dr*BphP_*dark*_ (blue) and *Dr*BphP_1*ps*_ (beige) structures around the A-ring, Asp207 and pyrrole water (PW). The structural model was inconclusive as to whether the A-ring twists around the double bond between the B- and A-ring, or whether it tilts downward hinged on the connection between C- and B-ring. (**b**). The regions of the pyrrole water (PW) and the area between the pyrrole rings show negative and positive density, respectively. The observed difference electron density is contoured at 3.3*σ*. (**c**). Density displayed for the backbone below the A-ring, including side chains of the strictly conserved Ile208 and Tyr176 as well as the surrounding water network. Monomer A is shown in this figure.

Furthermore, correlated negative and positive electron density features are observed on backbone atoms of the highly conserved stretch from Pro201 to His209. These difference electron density features indicate that the residues move away from the centre of the chromophore by 0.54 Å and 0.57 Å in monomers A and B, respectively (Supplementary Fig. 8b). The stretch includes Asp207 and it is located between the A-ring and the PHY domain, which makes it plausible that the changes in the chromophore cause this protein rearrangement. The changes are complemented by significant rearrangements of a stretch of waters and a conserved Tyr176 (Fig. 3c).

The photodissociation of the pyrrole water from the chromophore is a surprising finding. The pyrrole water is ubiquitously found in phytochrome structures (Burgie, Wang, et al. 2014; Burgie, Zhang, et al. 2016; Essen et al. 2008; Otero et al. 2016; Schmidt et al. 2018; Wagner, Brunzelle, et al. 2005; X. Yang et al. 2008; Xiaojing Yang et al. 2011). Our fluence dependent SFX data show that the negative density on the pyrrole water is the last signal to disappear when lowering the photon excitation densities 10-fold (Supplementary Fig. 3). This makes us confident that the photodissociation is in the single-photon absorption regime. The removal of the water requires significant energy, because hydrogen bonds to the A-, B-, and C-ring of the chromophore and the backbone C=O group of Asp207 have to be broken. We do not think that the twist of the D-ring causes this through direct steric interactions, because there is no contact between the pyrrole water and the D-ring. Rather, it may be triggered by an excited state charge redistribution between the pyrrole water and the chromophore, for example by ultrafast hydrogen or electron transfer (Toh et al. 2010). Such charge redistributions are typically facilitated by changes in geometry (Nosenko et al. 2008) and may therefore be caused indirectly by structural changes of the A-, C-, or D-rings, but this requires further investigation.

Conformational changes of the A-ring, Asp207 and the pyrole water have not been considered to occur on picosecond time scales. The strictly conserved Asp207 is a key residue for signal transduction, because it connects the chromophore to the PHY-tongue in Pr and Pfr (Essen et al. 2008; Takala et al. 2014; X. Yang et al. 2008). Its displacement suggests, together with the relocation of the residue stretch surrounding it, that disruption of the GAF-PHY interface occurs as early as 1 ps after photoexcitation (Supplementary Fig. 8b). With a hydrogen bond to the pyrrole water and in tight steric contact with the A-ring, Asp207 thereby acts as an extended arm of the chromophore. We propose that the photodissociation of the pyrrole water from the bilin and the change of the A-ring are integral parts of ultrafast phytochrome signalling towards the PHY domain.

## Discussion

Taken together, our data reveal a highly collective primary photoresponse for phytochromes. This is consistent with the fact that most point mutations of conserved residues alter, but do not inhibit, photoconversion (Wagner, Zhang, et al. 2008). The ultrafast structural changes are more extensive than in bacteriorhodopsin, photoactive yellow proteins, and in a fluorescent protein (Coquelle et al. 2018; Nogly et al. 2018; Pande et al. 2016). While previously observed ultrafast backbone movements have been interpreted as ‘protein quakes’ for myoglobin and bacteriorhodopsin (Barends et al. 2015; Nogly et al. 2018; Toh et al. 2010), the present backbone motion in the phytochrome binding pocket are much more directed (Supplementary Fig. 8). They occur in highly conserved regions of the protein and are part of the collective signalling response of the entire binding pocket.

We demonstrate that within one picosecond, a twist of the D-ring liberates the chromophore from the protein (Fig. 4a) and that movements of the pyrrole water, the A-ring and Asp207 lead to signalling directed towards the PHY-tongue (Fig. 4b). When mapped on the structure of the complete photosensory core module (Takala et al. 2014), both changes work together to destabilise the Arg466:Asp207 salt bridge. Tyr263 moves up, caused by the twist of the D-ring, and Asp207 moves down, caused by changes of the A-ring, retracting both residues from the salt bridge.

**Figure 4:**
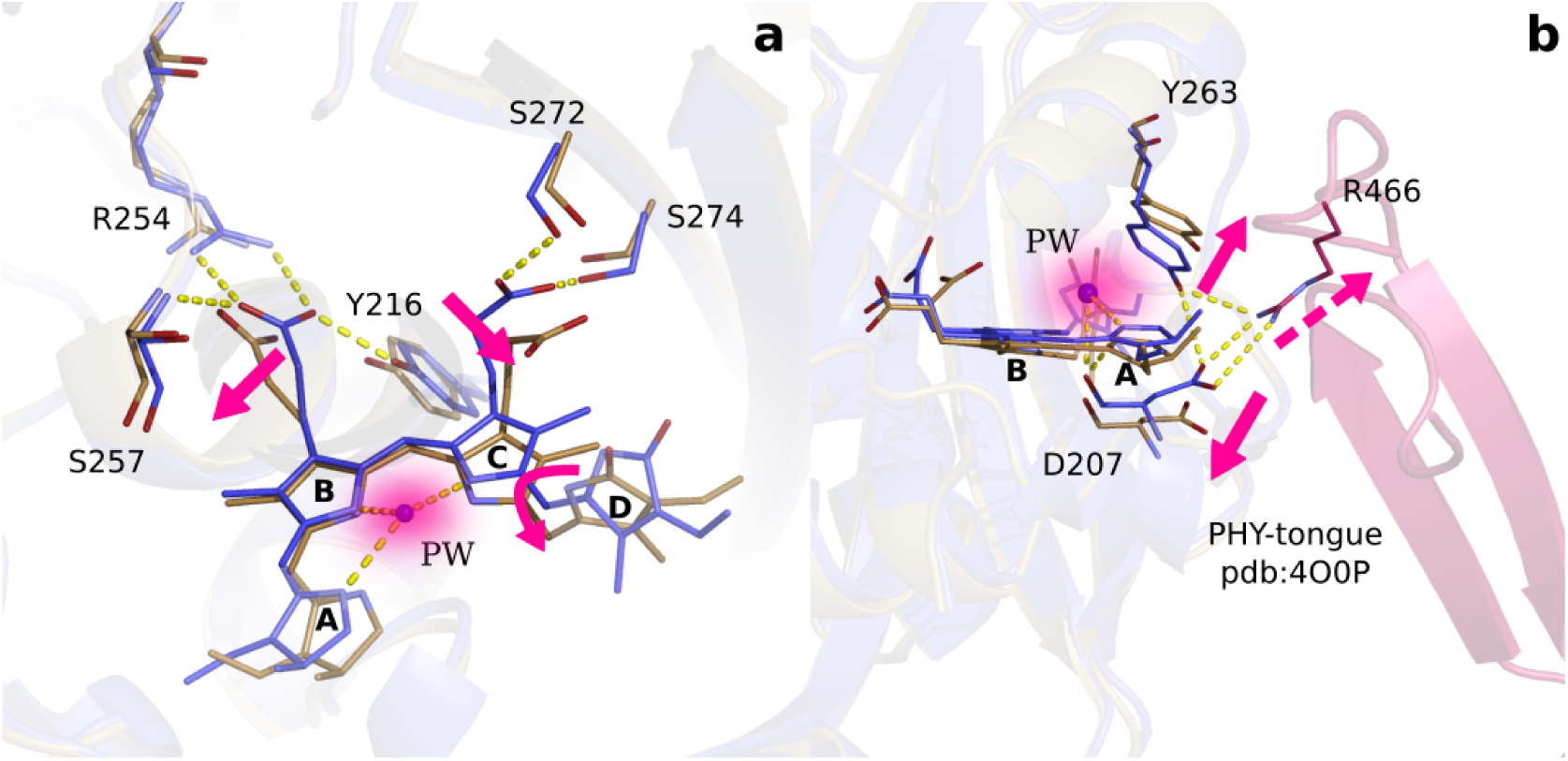
Two correlated photochemical events guide the primary photorespone of phytochrome proteins. (**a**). The structures (*Dr*BphP_*dark*_, blue and *Dr*BphP_1*ps*_, beige) indicate that rotation of the D-ring initiates breakage of non-covalent bonds of the propionates to the protein scaffold. Even the C- and A-rings are displaced significantly and the pyrrole water is dislocated from its original location at 1 ps (shade). (**b**). The same structures are overlayed with the complete photosensory core in Pr state (PDB ID 4O0P, pink) (Takala et al. 2014). The scissor-like separation of Asp207 and Tyr263 could result in breakage of the hydrogen bonds to Arg466 of the conserved PRXSF motif located in the PHY-tongue region.

Phytochromes have to be able to stabilise the bilin and to direct its photoisomerization from two photochemical ground states, Pr and Pfr. These differ both structurally and electronically, which precludes a single reaction trajectory for isomerization in the two directions. With this in mind, the observed primary photoresponse is reasonable. The structural signal is highly delocalized already at 1 ps, causing near-simultaneous liberation of the chromophore and initial signal transduction. We propose that these reaction trajectories stabilise each other, navigating the protein into a productive reaction path. The multidimensional reaction trajectory is consistent with the low quantum yields for photoconversion (Lamparter et al. 1997), which are characteristic for the phytochrome superfamily. Whereas the twisting motion of the D-ring has been the working model for phytochrome activation and is now confirmed, the photodissociation of the pyrrole water is highly surprising. We propose that both chemical events work together and enable phytochrome proteins to translate light information into structural signals, guiding the growth and development of bacteria, fungi, and plants on Earth.

## Materials and Methods

### Protein purification and crystallization

The *His*_6_-tagged PAS-GAF domain from *D*. *radiodurans* (aa 1-321) in vector pET21b(+) (Wagner, Brunzelle, et al. 2005) was expressed and purified like previously described (Lehtivuori et al. 2013; Takala et al. 2014). The recombinant protein was expressed in *Escherichia coli* strain BL21(DE3), either with or without *Ho*1 to yield holo- or apoprotein, respectively. Cells were lysed with Emulsiflex® and cleared by centrifugation (20 000 rpm, 30 min, +4°C). Full biliverdin incorporation was ensured by adding 8 mg of biliverdin hydrochloride per litre of cell culture (Frontier Scientific) to the cell lysate, followed by overnight incubation on ice. The protein was then purified at room temperature with HisTrap HP column (GE Healtcare) in 30 mM Tris, 50 mM NaCl and 5 mM imidazole (pH 8) and eluted with increasing imidazole concentration (gradient elution over 5-500 mM). Size-exclusion chromatography was then conducted with a HiLoad 26/600 Superdex 200 pg column (GE Healthcare) in buffer (30 mM Tris pH 8.0). Finally, the protein was concentrated to 30-50 mg/mL and flash-frozen in liquid nitrogen.

Crystals were set up under green safe light and grown in dark. Batch crystallization was performed as described (Edlund et al. 2016). 50 *µ*L of purified protein (25-30 mg/mL) was added to 450 *µ*L of reservoir solution (60 mM Sodium acetate pH 4.95, 3.3% PEG 400, 1 mM DTT and 30% 2-methyl-2,4-pentanediol) and immediately mixed. Initial microcrystals were grown on a tipping table at 4 °C for 48h. Once the microcrystals were formed, additional protein was added to increase crystal size. The microcrystals were first pelleted by brief centrifugation and 400 *µ*L of supernatant was removed. 200 *µ*L of diluted protein (14 mg/mL in 30 mM Tris pH 8.0) was then added to the microcrystals along with 200 *µ*L of fresh reservoir solution. After 48h incubation on a tipping table at room temperature, crystals of diffraction quality (20-70 *µ*m long needles) were formed (Supplementary Fig. **2**).

### Light scattering of the grease jet

The visible absorption spectra of the grease, grease mixed with crystallisation buffer, microcrystals in the grease matrix, resembling the characteristics of the jet during the XFEL experiments, were measured. 2.5 *µ*l of sample was placed between two CaF_2_ windows with a 50 *µ*m Teflon spacer and squeezed together. The UV-Vis spectra were measured with a transmittance diode-array UV-Vis spectrometer (Cary 8454, Agilent Technologies). Despite low absorption in the visible spectral range, the grease (Superlube, Synchochemical corp.) absorption properties changed drastically when mixed with crystallisation buffer, showing a strong scattering effect. The absorbance of the microcrystals in the grease matrix was found to be OD *>*2 at 640 nm with a pathlength of 50 *µ*m (Supplementary Fig. **1**c). Thus the number of photons that hit the crystals in the 100 *µ*m or 75 *µ*m jet, which was the path length used at the XFEL experiment, is significantly reduced due to the scattering loss. The crystals see only a fraction of the incoming light intensity, similar to what Nogly *et al*. estimated (Nogly et al. 2018). The raw spectra of micorcrystals (Supplementary Fig. **1**d) in the buffer has been measured between two CaF_2_ windows without spacer to minimise the absorption loss, the pathlength was estimated to be ≤ 50 *µ*m.

### SFX Data acquisition

Serial femtosecond crystallographic data were collected at SPring-8 Angstrom Compact Free Electron Laser (SACLA) in two beamtimes in October 2018 and May 2019. The microcrystals were pelleted by brief centrifugation, the crystal pellet was mixed with 180 *µ*L of grease, and loaded into a 4 mm sample reservoir for data acquisition. The sample was delivered to the X-ray beam at a flow rate of 2.5 *µ*L/min or 4.2 *µ*L/min for 75 *µ*m and 100 *µ*m diameter nozzles, respectively. The time resolution of the experiment was limited by the jitter of the XFEL of 100 fs *r*.*m*.*s*..

### Data processing

The background of the detector was estimated by averaging the first 150 dark images in each run and then subtracted from each diffraction pattern. Diffraction images with Bragg spots (the “hits”) were found by a version of Cheetah adapted for SACLA (Barty et al. 2014; Nakane et al. 2016). These hits were indexed by the program CrystFEL (version 0.6.3) (White et al. 2012). Indexing was performed using Dirax and Mosflm (Battye et al. 2011; Duisenberg and IUCr 1992). Spot finding in each diffraction image was done with the peakfinder8 algorithm using the parameters (min SNR = 4.5, threshold = 100, minimum pixel counts = 3). The indexed patterns were merged and scaled using partialator in CrystFEL and hkl files were produced. The figure of merits (Supplementary Table 1) were calculated by using compare hkl and check hkl in CrystFEL.

### Refinement of dark structure

The initial phases were solved by molecular replacement with Phaser (McCoy et al. 2007) and the PAS-GAF crystal structure (PDB ID 5K5B) (Edlund et al. 2016) as a search model. The structure was refined with REFMAC version 5.8.0135 (Murshudov et al. 2011) with a weight factor for the geometry restraints of 0.05, accompanied by model building steps with Coot 0.8.2 (Emsley et al. 2010). The final structure (*Dr*BphP_*dark*_) had Rwork/Rfree of 0.161/0.192 and no Ramachandran outliers (Supplementary Table 1).

### Computation of difference electron density maps

The difference structure factor (Δ*F*) were computed from the measured structure factor amplitudes in dark and for preset delay times between laser and X-ray pulses as 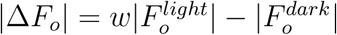 and with phases taken from the dark structural model (*Dr*BphP_*dark*_). 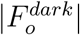 and 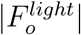 were brought to the absolute scale by first scaling 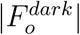 to 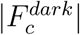 and then scaling 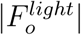 to 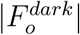 using the CCP4 program Scaleit (Winn et al. 2011). A weighting factor (*w*) was determined for each reflection to reduce the influence of outliers (Ren et al. 2001). From the weighted Δ*F*, a difference electron density map (Δ*ρ*) is calculated using the program ‘fft’ from the ccp4 suite of programs (Winn et al. 2011). Since Δ*F* are on the absolute scale, Δ*ρ* is on half the absolute scale as a result of the difference Fourier approximation.

### Structure refinement of light structure

Extrapolated structural factors were assembled from amplitudes computed as 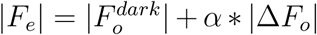, where 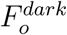 are the amplitudes measured in dark and *α* is inversely related to the population of the photoinduced state, phases were taken from *Dr*BphP_*dark*_. *F*_*e*_ represents the pure structure factor of the excited state (Supplementary Fig. 5). Refinement of structural models was then performed in real and reciprocal space, using Coot (Emsley et al. 2010) and Phenix (Adams et al. 2010). The equilibrium values for the restraints used in the refinement of the biliverdin chromophore were taken from a minimal energy biliverdin ground state geometry that was obtained at the B3LYP/6-31G* level of density functional theory. Torsional restraints for the excited state geometry with the twisted D-ring were obtained at the SA(5)-CASSCF(12,12)/cc-pVDZ level of ab initio theory. We removed the torsion restrains for the C/D-ring (C14-C15-C16-C17; C14-C15-C16-ND; C13-C14-C15-C16; NC-C14-C15-C16) and for the A/B-ring (C3-C4-C5-C6; NA-C4-C5-C6; C4-C5-C6-C7; C4-C5-C6-NB) during refinement. The overall aim of the refinement was to maximise the agreement between the observed and calculated difference maps. To evaluate the agreement, we subtracted the calculated from the observed difference electron density (Δ*ρ*_*o*_ − Δ*ρ*_*c*_) and used the resulting map to objectively increase correspondence throughout refinement steps. To compute the difference-difference maps, the highest and lowest intensities of Δ*ρ*_*o*_ were scaled to the corresponding maximum and minimum of Δ*ρ*_*c*_ and the observed Δ*ρ*_*o*_ were interpolated linearly according to this scaling. Calculation of Pearson Correlation Coefficient (PCC) values between the Δ*ρ*_*o*_ and Δ*ρ*_*c*_ were applied to guide refinement of specific regions, such as the D-ring and the whole chromophore region. To do so, the correlation was determined based on electron density within a sphere with a radius of 3.5 Å or 10 Å centred on the D-ring or pyrrole water, respectively. As a final step in the refinement procedure, we refined the models with REFMAC version 5.8.0135 (Murshudov et al. 2011) with high geometry restraints (weight factor 0.005). This was done against phased extrapolated structure factors, using the phases of the refined light and dark structure for computation of phased Δ*F* as described (Pande et al. 2016). The structures did not change much, although the R factors dropped in this last step of refinement to Rwork/Rfree of 0.230/0.256 (Supplementary Table. 1).

## Acknowledgments

The experiments at SACLA were performed at BL3 with the approval of the Japan Synchrotron Radiation Research Institute (JASRI) (Proposal No. 2018A8055 and 2019A8007). S.W. acknowledges the European Research Council for support (grant number: 279944). This work was supported by Academy of Finland grants 285461 and 296135 (H. T. and J.A.I., respectively) and Jane and Aatos Erkko foundation (J.A.I.). This research is partially supported by Platform Project for Supporting Drug Discovery and Life Science Research (Basis for Supporting Innovative Drug Discovery and Life Science Research (BINDS)) from Japan Agency for Medical Research and Development (AMED). We thank Dr. Takanori Nakane for assistance with data processing during the beamtime. This work was supported by NSF Science and Technology Centers grant NSF-1231306 (“Biology with X-ray Lasers”). This work has been done as part of the BioExcel CoE (www.bioexcel.eu), a project funded by the European Union contracts H2020-INFRAEDI-02-2018-823830 and H2020-EINFRA-2015-1-675728. S.W., M.S., K.M., and J.A.I. conceived the experiments S.W. and E.C. planned them, and E.C., W.Y.W., H.T., and L.C. purified the protein and prepared the microcrystals. E.C., W.Y.W., H.T., S.P., L.C., V.K., L.H., M.C., M.P., J.K., R.N., L.I., A.N., A.C., M.M., R.B., E.N., R.T., T.T., S.I., L.F., G.G., E.S., J.A., M.S., and S.W. acquired the data at SACLA. V.K. and M.K. measured the UV-Vis spectroscopic data. S.P. processed the crystallography data and E.C., W.Y.W., H.T., M.S. and S.W. computed difference electron density maps and refined the structural models. G.G. and D.M. provided the geometry restrains biliverdin in the excited state. S.W., M.S. and E.C. wrote the paper with input from all authors.

---

**Table 1:** Crystallographic table.

|  | **Dark** | **1 ps** | **10 ps** |
| --- | --- | --- | --- |
| **PDB code** | TBD | TBD |  |
| **Data collection** |  |  |  |
| Temperature (K) | 293 | 293 | 293 |
| Space Group | P212121 | P212121 | P212121 |
| **Cell dimensions (a, b, c)** |  |  |  |
| a, b, c (Å) | 54.98 116.69 117.86 | 54.98 116.69 117.86 | 54.98 116.69 117.86 |
| $\alpha, \beta, \gamma$ (°) | 90.0 90.0 90.0 | 90.0 90.0 90.0 | 90.0 90.0 90.0 |
| Data resolution overall (Å)<sup>‡</sup> | 45.77-2.07<br>(2.10-2.07) | 41.46-2.21<br>(2.25-2.21) | 45.77-2.14<br>(2.17-2.14) |
| $R_{split}$ (%)<sup>‡</sup> | 5.79 (120.05) | 10.59 (114.64) | 5.70 (121.86) |
| SNR ($I/\sigma(I)$)<sup>‡</sup> | 9.21 (0.83) | 6.10 (0.88) | 10.11 (0.99) |
| CC(1/2)<sup>‡</sup> | 0.99 (0.33) | 0.98 (0.38) | 0.99 (0.344) |
| Completeness (%)<sup>‡</sup> | 100 (100) | 100 (100) | 100 (100) |
| Multiplicity<sup>‡</sup> | 461.35 (65.9) | 106.11 (34.3) | 347.36 (62.1) |
| Number of hits | 149074 | 42853 | 159997 |
| Number of indexed patterns | 70726 | 21150 | 70335 |
| Indexing rate(%)<sup>§</sup> | 47.44 | 49.35 | 43.96 |
| Number of total reflections | 24017763 | 5310179 | 17823530 |
| Number of unique reflections | 52060 | 39316 | 43279 |
| **Refinement** |  |  |  |
| Resolution (Å)<sup>‡</sup> | 45.82-2.07<br>(2.12-2.07) | 36.94-2.21<br>(2.27-2.21) |  |
| $R_{work} / R_{free}^{\dagger}$ | 0.162/0.191<br>(0.317/ 0.346) | 0.230/0.256<br>(0.411/0.443) | |
| Number of atoms | 5123 | 5135 |  |
| Average B factor (Å<sup>2</sup>) | 76.44 | 78.63 |  |
| **R.m.s deviations** |  |  |  |
| Bond lengths (Å) | 0.007 | 0.006 |  |
| Bond angles (°) | 1.251 | 1.152 |  |

<sup>‡</sup> Highest resolution shell is shown in parentheses.
<sup>§</sup> Ratio of the number of indexed images to the total number of hits.

**Table 2:** Difference electron density features listed for certain atoms.

| Object | Label | 1ps<br>A | 1ps<br>B | 10ps<br>A | 10ps<br>B |
| --- | --- | --- | --- | --- | --- |
| **Pyrrole Water** |  |  |  |  |  |
| Pyrrole Water (-) | **XV** | -8.2σ | -9.4σ | -5.0σ | -6.5σ |
| Pyrrole Water (+) Alt. 1 | **XVI** | 4.8σ | 4.2σ | 3.0σ | 3.4σ |
| Pyrrole Water (+) Alt. 2 | **XVII** | 4.4σ | 5.8σ | 3.0σ | 3.3σ |
| **D-ring** |  |  |  |  |  |
| D-ring N-H/C=O (-) | **I** | -4.2σ | -7.3σ | -6.4σ | -5.1σ |
| D-ring Methyl (-) | **II** | -4.3σ | -3.5σ | -2.9σ | -4.9σ |
| D-ring Vinyl (-) | **III** | -4.0σ | -4.8σ | -3.5σ | -3.4σ |
| D-ring N-H/C=O (+) | **IV** | 4.6σ | 6.4σ | 4.6σ | 4.1σ |
| D-ring Methyl (+) | **V** | 3.5σ | 3.4σ | - | - |
| D-ring Vinyl (+) | **VI** | 4.1σ | 4.6σ | 3.4σ | 2.6σ |
| **C-ring** |  |  |  |  |  |
| C-propionate (-) | **IX** | -6.4σ | -5.8σ | -5.9σ | -4.7σ |
| C-ring (-) | **VII** | -5.5σ | -4.4σ | -5.1σ | -3.2σ |
| C-propionate (+) | **X** | 6.9σ | 6.4σ | 7.4σ | 4.9σ |
| C-ring (+) | **VIII** | 4.9σ | 5.2σ | 3.2σ | 5.4σ |
| **A-ring** |  |  |  |  |  |
| A-ring C=O (-) | **XVIII** | -4.3σ | -4.7σ | -3.6σ | -2.9σ |
| A-ring C=O (+) | **XIX** | 5.2σ | 5.0σ | 5.7σ | 4.7σ |
| **His260** |  |  |  |  |  |
| His260 Sidechain(-) | **XI** | -6.7σ | -5.2σ | -6.3σ | -4.1σ |
| His260 Sidechain(+) | **XII** | 4.3σ | 4.5σ | 3.5σ | 3.2σ |
| **Tyr263** |  |  |  |  |  |
| Tyr263 Sidechain(-) | **XIII** | -6.9σ | -7.0σ | -4.7σ | -6.2σ |
| Tyr263 Sidechain(+) | **XIV** | 4.8σ | 4.9σ | 3.0σ | 4.3σ |
| **Asp207** |  |  |  |  |  |
| Asp207 Sidechain (-) | **XX** | -6.2σ | -6.7σ | -5.6σ | -3.4σ |
| Asp207 Sidechain(+) | **XXI** | 5.7σ | 5.0σ | 4.2σ | - |
| Asp207 Backbone (-) | **XXII** | -6.2σ | -5.4σ | -5.1σ | -4.5σ |
| Asp207 Backbone (+) | **XIII** | 4.6σ | 4.4σ | 4.5σ | 3.5σ |

**Supplementary Figure 1:**
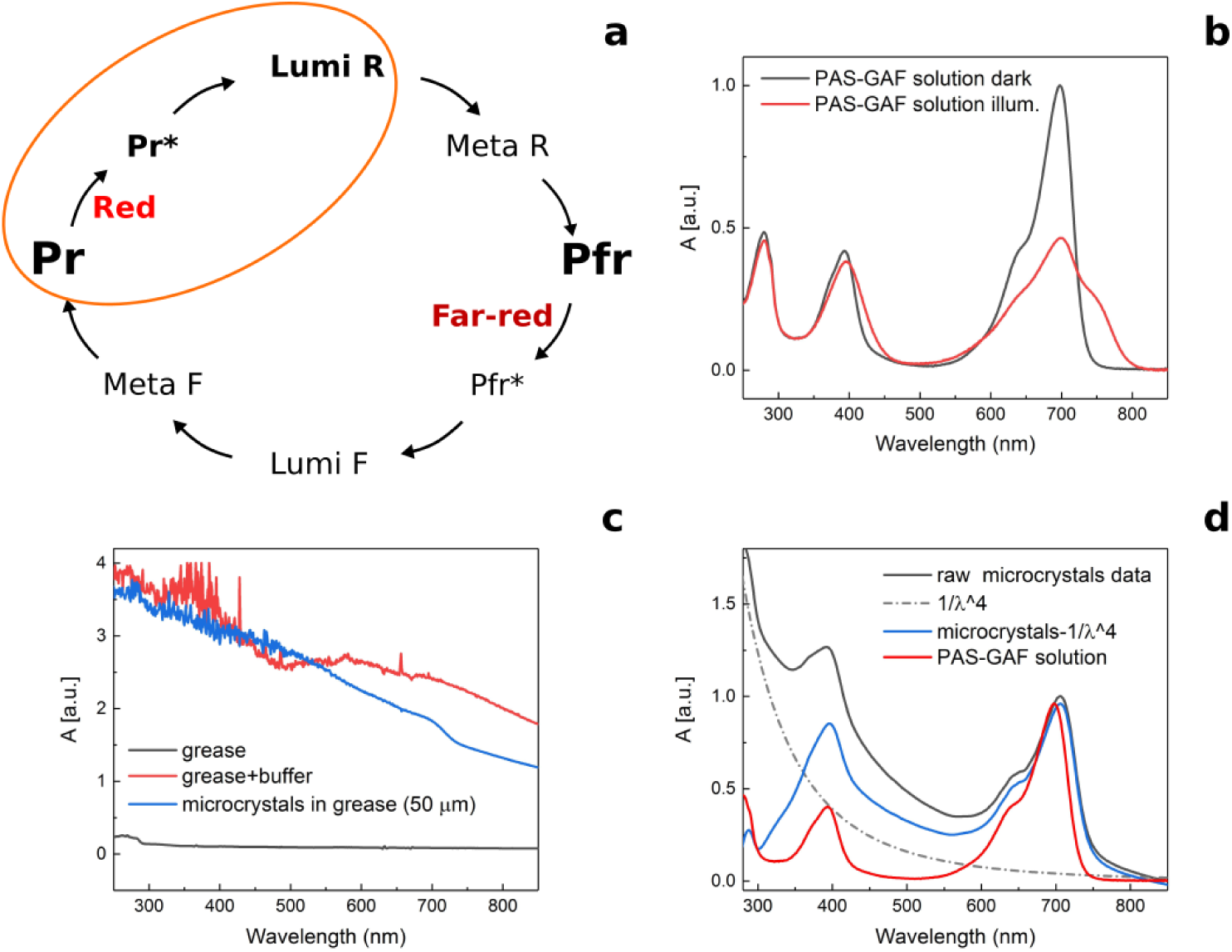
The photocycle of *Deinococcus radiodurans* phytochrome (*Dr*BphP) and the spectral properties of its PAS-GAF fragment. (**a**). Schematic illustration of the photoconversion between Pr and Pfr states in *Dr*BphP. We note that the final state of the PAS-GAF fragment (also referred to as chromophore-binding domain, CBD) does not have the same spectra shape as in full-length phytochromes. (**b**). Absorption spectra of the dark (Pr) and illuminated (Pfr) *Dr*BphP_*CBD*_ fragments in solution, indicating functionality of the protein. (**c**). Absorption spectra of the grease (black), grease mixed with crystallisation buffer (red), and *Dr*BphP_*CBD*_ microcrystals in the grease matrix (measured in the 50 *µ*m path length). Significant (OD *>*2) light scattering occurs when the grease is mixed with the crystal buffer. The scattering varied with the mixing procedure, but we estimate that the light intensities inside the grease jet are attenuated by at least one order of magnitude. (**d**). Raw absorption spectra of the *Dr*BphP_*CBD*_ microcrystals in buffer (black), scattering correction (1/*λ*^4^, dash dot), scattering corrected absorption spectra of the *Dr*BphP_*CBD*_ microcrystals (blue), and *Dr*BphP_*CBD*_ in solution (red).

**Supplementary Figure 2:**
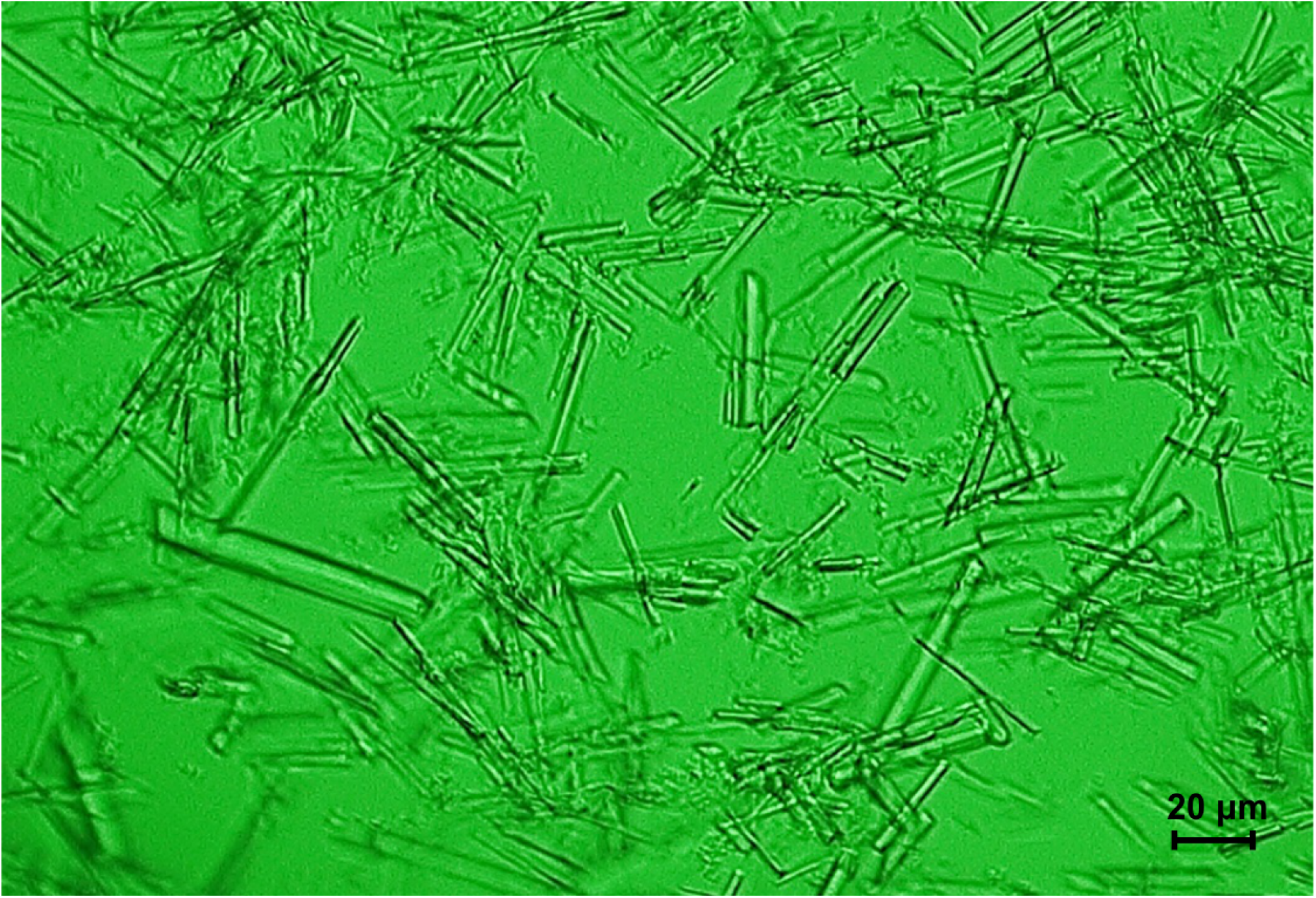
Microcrystals used for SFX data acquisition in SACLA. The crystals reported here were shaped as needles of 20-70 *µ*m.

**Supplementary Figure 3:**
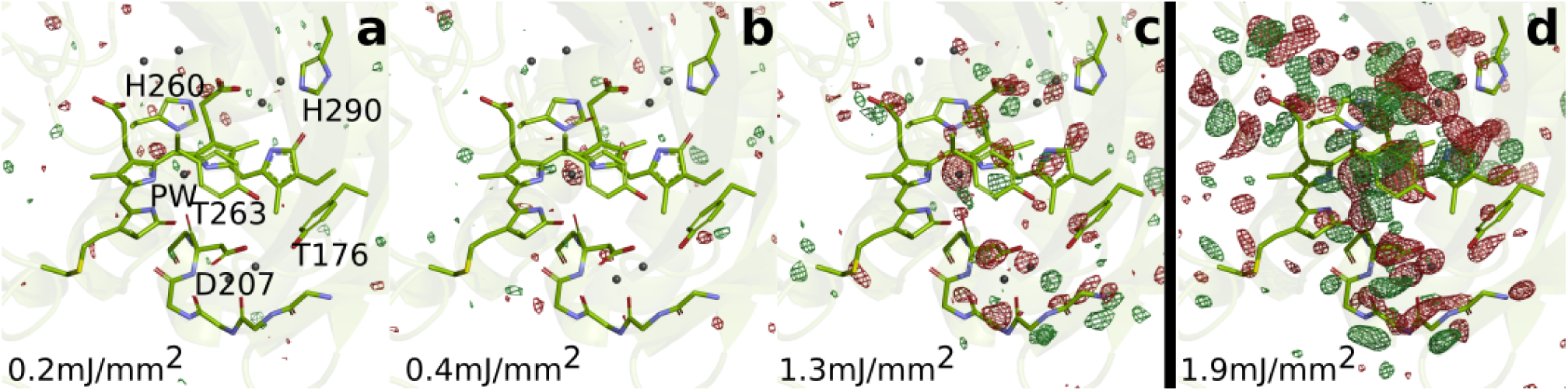
SFX data as a function of excitation fluence. The *Dr*BphP_*dark*_structure (green) is shown together with the observed difference electron density, contoured at 3.5, at 1 ps collected using (**a**) 0.2 *µ*J/mm^2^, (**b**) 0.4 *µ*J/mm^2^, (**c**) 1.3 *µ*J/mm^2^, and (**d**) 1.9 *µ*J/mm^2^. All spot sizes were computed assuming Gaussian line shapes with the (1*/e*^2^) convention. The data shown in panel A-C were collected at SACLA in May 2019 whereas the data shown in panel D was collected in October 2018. The same experimental setup was used in both occasions. In the latter experiment the femtosecond laser beam was aligned to 40-50 *µ*m distance from the interaction spot between X-rays and jet in the direction of flow. The laser intensities were corrected for this displacement assuming a Gaussian line shape. The excitation fluence is similar to previous femtosecond time-resolved SFX experiments (Barends et al. 2015; Coquelle et al. 2018; Nogly et al. 2018; Pande et al. 2016), but higher than for phytochromes in solution. We attribute this to the high optical scattering cross section of the grease/buffer mixture (Supplementary Fig. **1**). Since the crystallographic signals were reduced when lowering the excitation fluence and disappeared completely when reaching 1/10 of the maximum value, we conclude that the excitation fluence that actually reaches the crystals in the grease matrix is much lower than the incoming photon fluence and that the photoexcitation is in the single-photon regime.

**Supplementary Figure 4:**
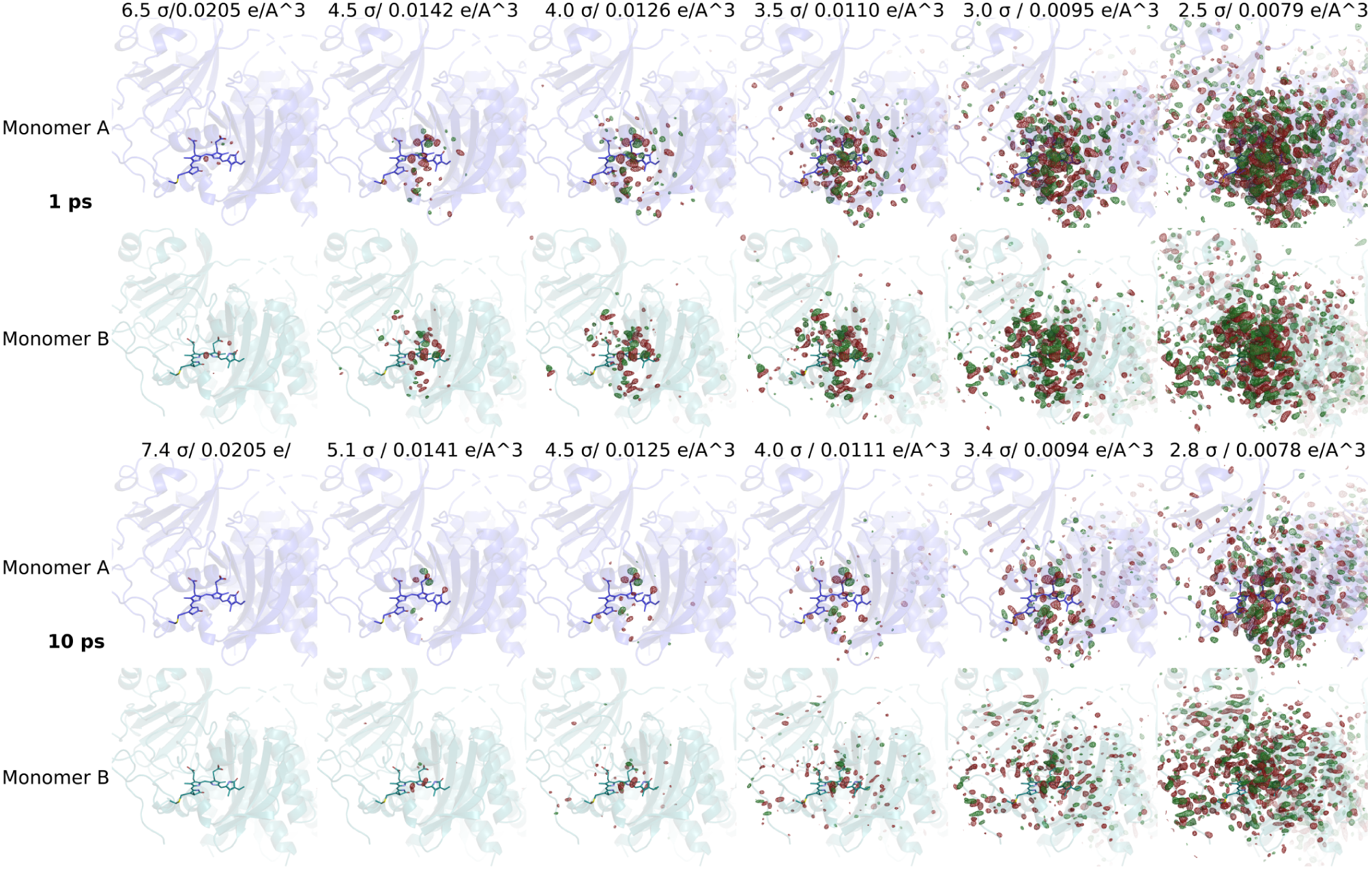
Significant difference electron density features observed. Difference electron density maps at 1 ps and 10 ps for monomer A and B are shown together with the *Dr*BphP_*dark*_ structure (monomer A: blue, monomer B: aqua). Red and green contour surfaces denote negative and positive densities, respectively. The maps are contoured at electron density levels as indicated. We determine the background level to be at 3*σ* for 1 ps and at 3.4*σ* for 10 ps, since below these levels random signals outside the chromophore region appear.

**Supplementary Figure 5:**
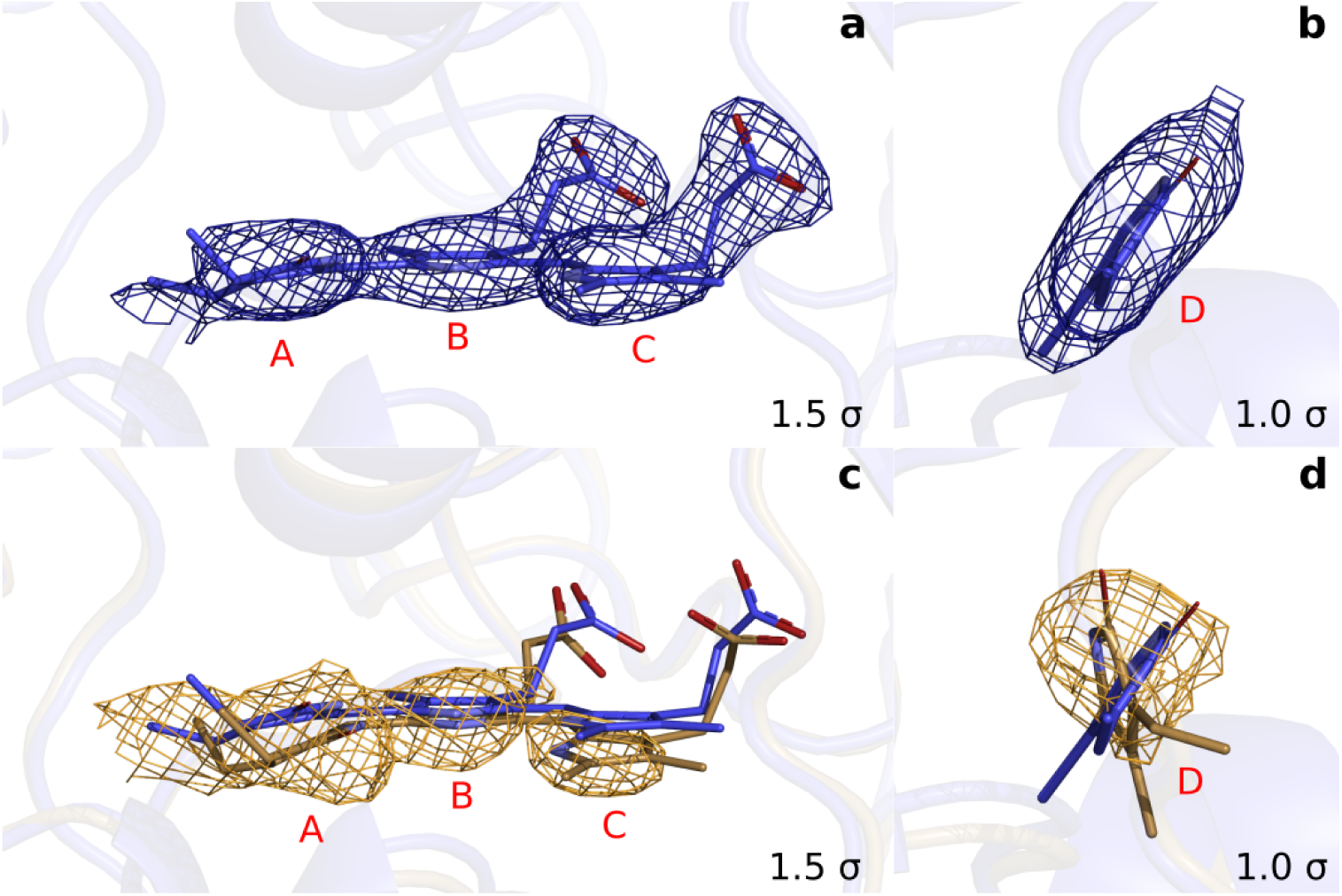
Extrapolated maps at *α* = 25 demonstrate that the D-ring twists in the excited state. (**a**). Rings A to C of monomer A together with the 2*F*_*o*_− *F*_*c*_ maps of the refined dark structure (*Dr*BphP_*dark*_: blue). (**b**). The same data for the D-ring in a different orientation. (**c**) and (**d**). Equivalently, the refined structure (*Dr*BphP_1*ps*_: beige) of the chromophore in monomer A together with the extrapolated map *FT* (*Fe*) at 1 ps. (*Fo*, phases from the dark structure). We observed a round-shape feature for the D-ring, but flat densities for C-ring (downward shift) and B-ring (almost unshifted). This suggests that the D-ring twists significantly. For the A-ring we observe a broad density, with the strongest contribution indicating a twisted or tilted ring.

**Supplementary Figure 6:**
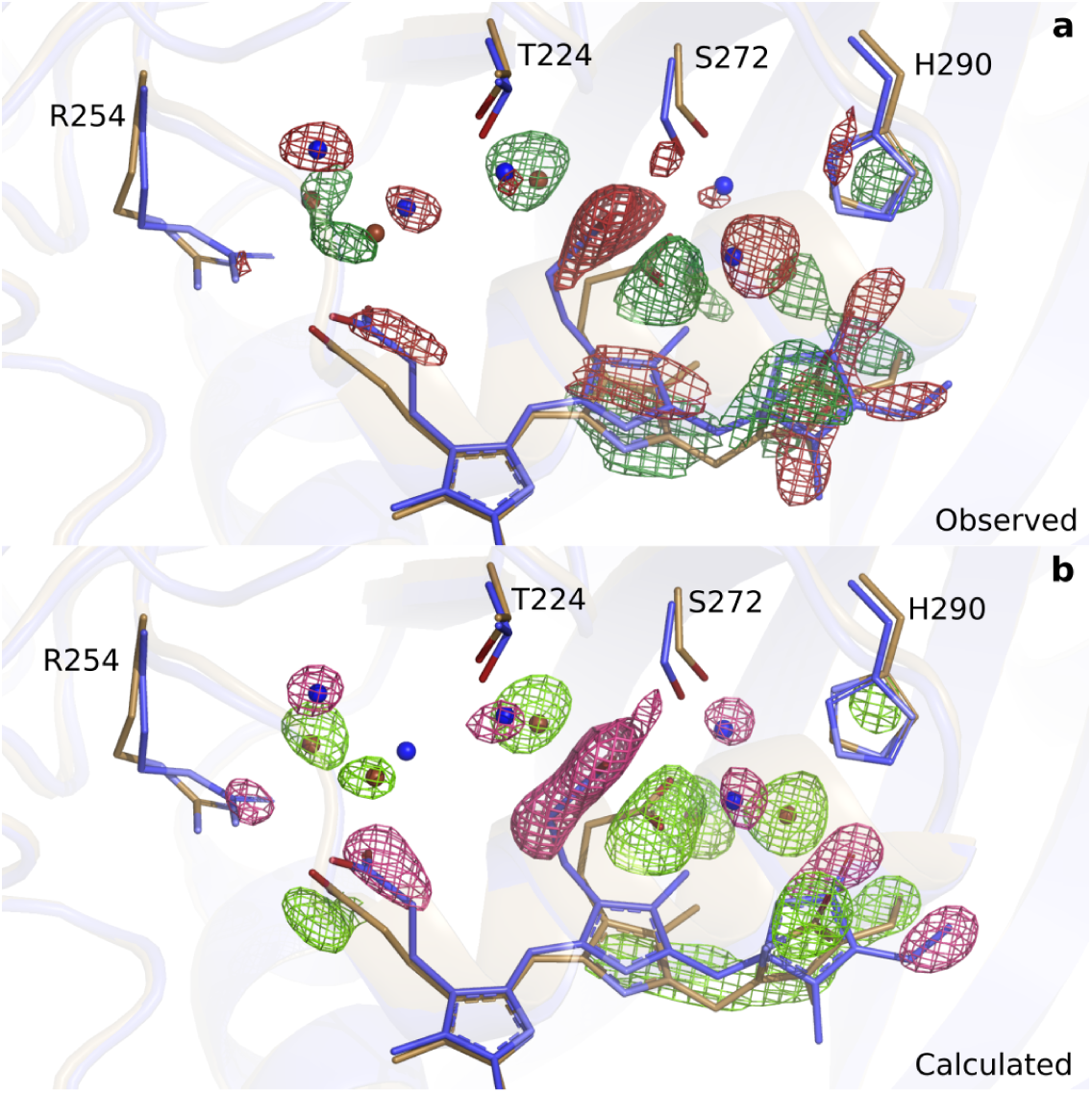
Comparison of observed (upper panel) and calculated (lower panel) difference electron densities indicate good agreement around the B-D rings. (**a**). The observed difference electron density map (3.3 *σ*) is displayed for the B-D ring surroundings of the *Dr*BphP_*dark*_(blue) and the *Dr*BphP_1*ps*_structure (beige) for monomer A. (**b**). The same view and structures displayed with the maps contoured at 4.5 *σ*.

**Supplementary Figure 7:**
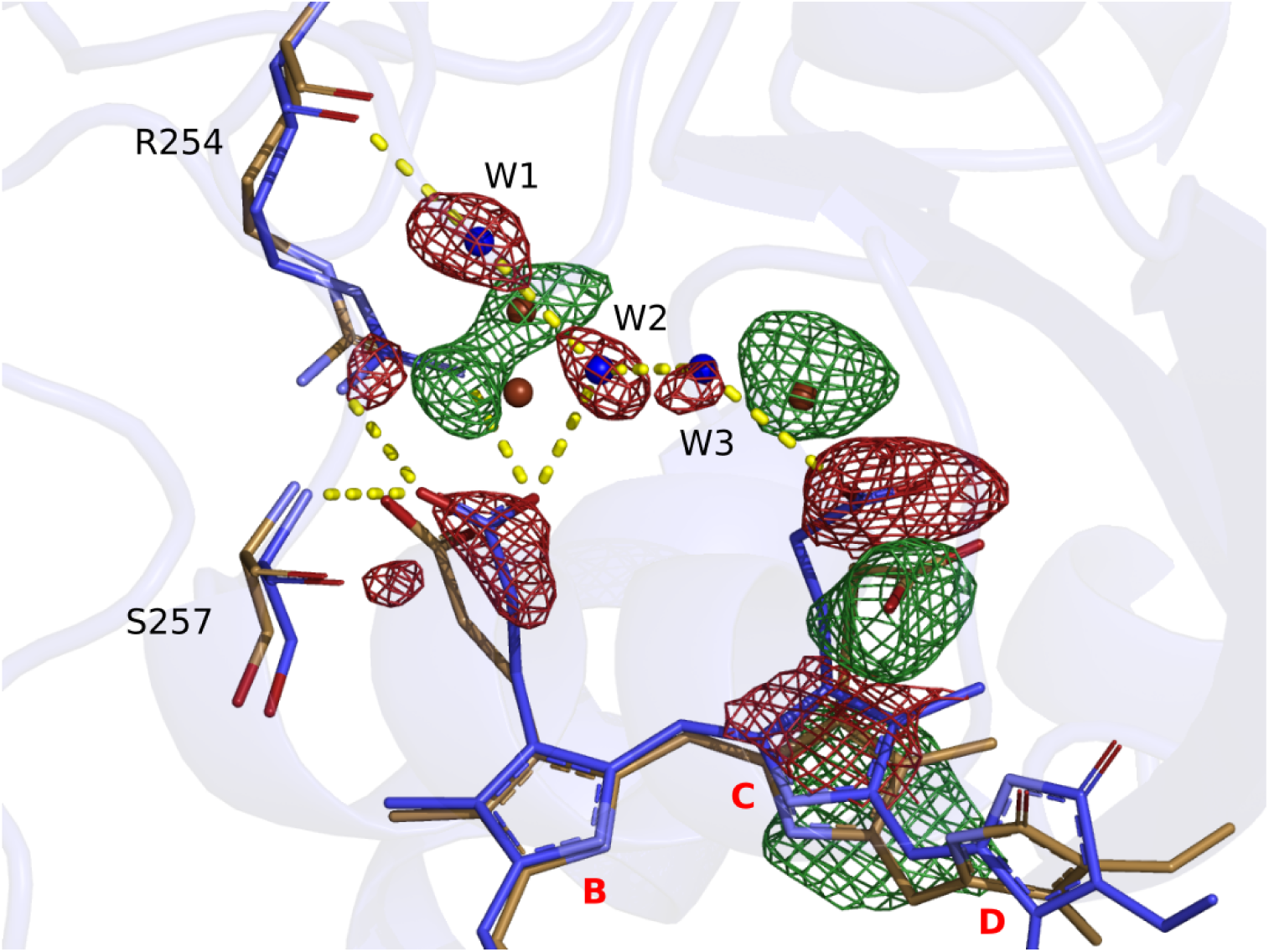
Hydrogen bonding network between the chromophore propionate groups is disrupted at 1 ps delay time. Correlating negative and positive density features indicate that the C-ring moves downwards, together with the C-ring propionate. The negative density features located on the propionate groups, three waters (W1, W2 and W3), Arg254, and Ser257 collectively indicate that the hydrogen bonding network is resolved at 1ps. Difference electron density features are not observed on the B-ring. *Dr*BphP_*dark*_is shown in blue and *Dr*BphP_1*ps*_in beige. The chromophore is shown for monomer A together with the observed difference map at 1 ps delay time, contoured at 3.0*σ*.

**Supplementary Figure 8:**
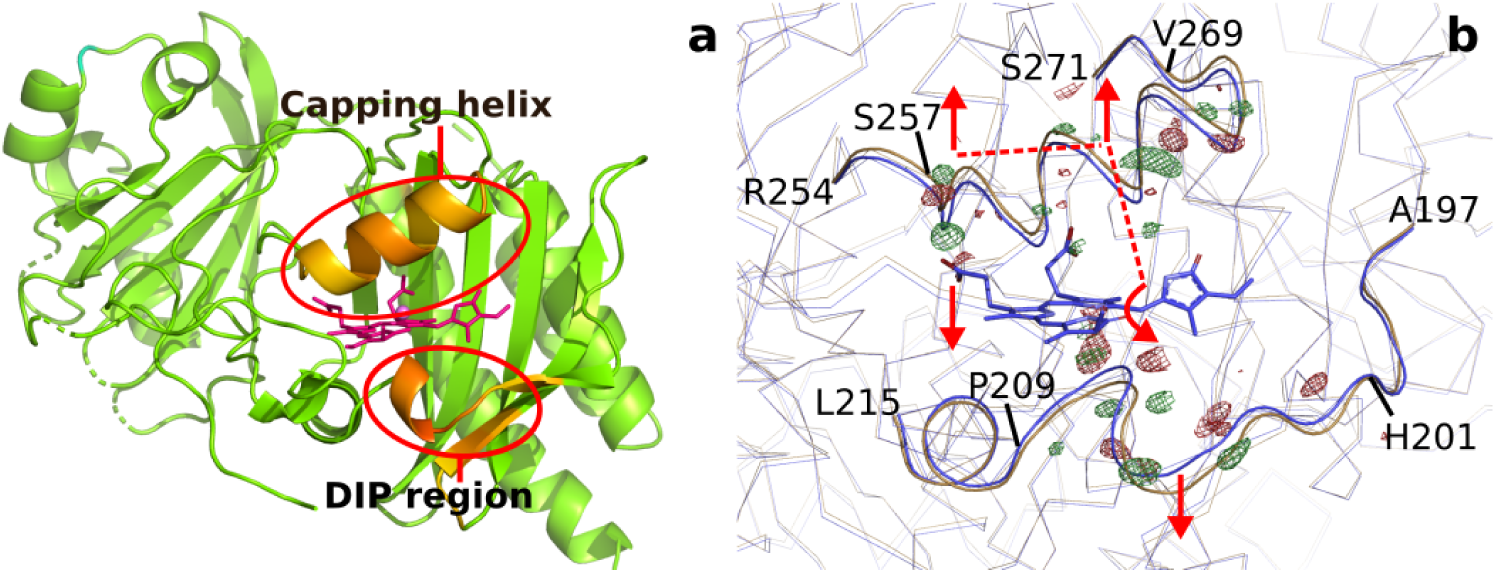
Backbone movements at 1 ps. (**a**). Difference (*Dr*BphP_1*ps*_-*Dr*BphP_*dark*_) in distances between each C*α* and the pyrrole water (*Dr*BphP_*dark*_) plotted for all differences that are larger than 0.45 Å for monomer B. Longer distances are coloured red and biliverdin is shown as pink sticks. The regions that move the most, the ‘DIP region’ and the ‘capping helix’ above the chromophore are marked with red circles. (**b**). The observed difference electron densities indicate backbone movements away from the chromophore for residues 201-209 and 257-269, and an expansion of the chromopore-binding pocket. The *Dr*BphP_*dark*_ structure is coloured blue and the *Dr*BphP_1*ps*_ structure is coloured beige. The observed electron density map is contoured at 4.0 *σ*. The dashed lines indicate the proposed signal transduction pathway from D-ring to B-ring propionate.

**Supplementary Figure 9:**
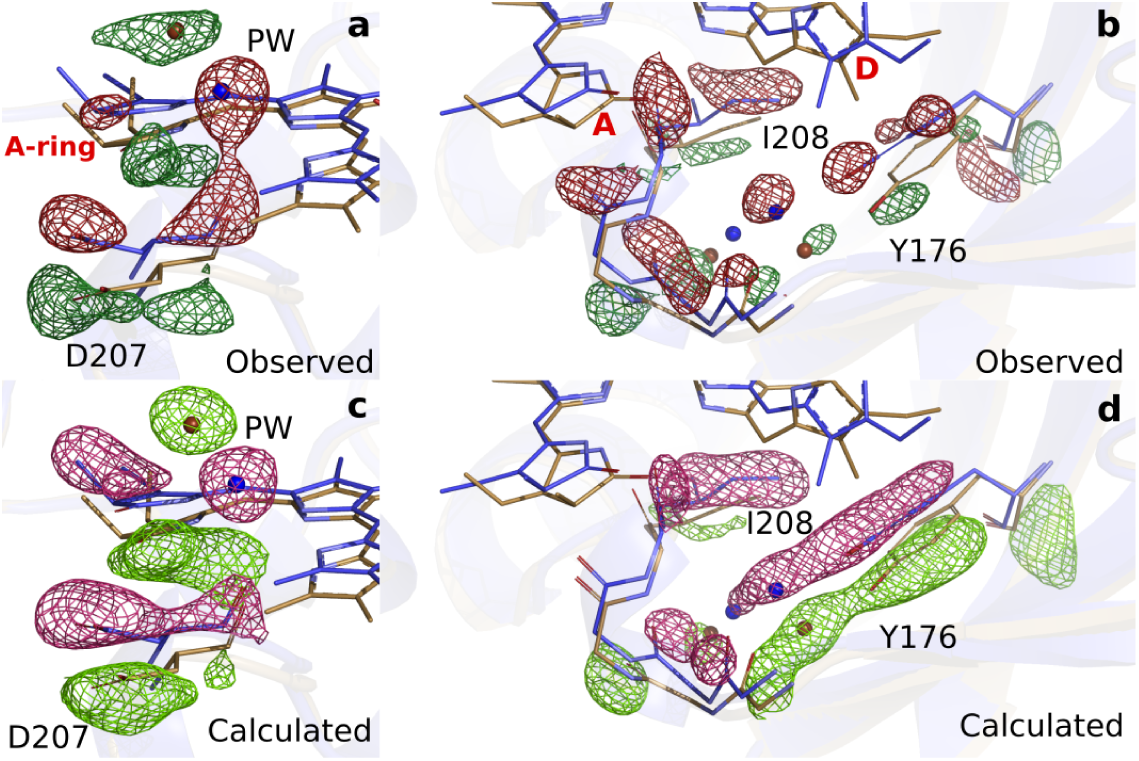
Comparison of observed and calculated difference electron densities in the A-ring region. (**a**). The observed difference electron density is displayed together with the *Dr*BphP_*dark*_(blue) and the *Dr*BphP_1*ps*_(beige) structures for monomer A. The data is shown around the A-ring, Asp207 and pyrrole water (PW). (**b**). For the backbone below the A-ring, side chains shown for the strictly conserved Ile208 and Tyr176 as well as the surrounding water network. (**c**). The same view as panel A displayed together with the calculated difference electron density. (**d**). The same view as panel B displayed together with the calculated difference electron density. The observed and calculated difference electron density maps are contoured at 3.3 *σ* and 3.5 *σ*, respectively.

